# Enhanced genome annotation strategy provides novel insights on the phylogeny of *Flaviviridae*

**DOI:** 10.1101/674333

**Authors:** Adriano de Bernardi Schneider, Denis Jacob Machado, Daniel Janies

**Affiliations:** AntiViral Research Center, Department of Medicine, University of California San Diego, San Diego, California, USA; Department of Bioinformatics and Genomics, College of Computing and Informatics, University of North Carolina at Charlotte, Charlotte, North Carolina, USA

## Abstract

The ongoing and severe public health threat of viruses of the family *Flaviviridae*, including dengue, hepatitis C, West Nile, yellow fever, and zika, demand a greater understanding of how these viruses evolve, emerge and spread in order to respond. Central to this understanding is an updated phylogeny of the entire family. Unfortunately, most cladograms of *Flaviviridae* focus on specific lineages, ignore outgroups, and rely on midpoint rooting, hampering their ability to test ingroup monophyly and estimate ingroup relationships. This problem is partly due to the lack of fully annotated genomes of *Flaviviridae*, which has genera with slightly different gene content, hindering genome analysis without partitioning. To tackle these problems, we developed an annotation pipeline for *Flaviviridae* that uses a combination of *ab initio* and homology-based strategies. The pipeline recovered 100% of the genes in reference genomes and annotated over 97% of the expected genes in the remaining non curated sequences. We further demonstrate that the combined analysis of genomes of all genera of *Flaviviridae* (*Flavivirus, Hepacivirus, Pegivirus*, and *Pestivirus*), as made possible by our annotation strategy, enhances the phylogenetic analyses of these viruses for all optimality criteria that we tested (parsimony, maximum likelihood, and posterior probability). The final tree sheds light on the phylogenetic relationship of viruses that are divergent from most *Flaviviridae* and should be reclassified, especially the soybean cyst nematode virus 5 (SbCNV-5) and the Tamana bat virus. We also corroborate the close phylogenetic relationship of dengue and zika viruses with an unprecedented degree of support.

## Introduction

The family *Flaviviridae* comprises the genera *Flavivirus, Hepacivirus, Pegivirus*, and *Pestivirus*, all of which share structural and genomic similarity (1). There are several examples that demonstrate the clinical relevance and zoonotic nature of this family. In humans, hepatitis C virus (HCV) causes a disease that damages the liver (2). Human pegivirus-1 (HPgV-1), causes human encephalitis (3) as well bovine viral diarrhea (BVDV), long-known cause of major losses in livestock (4). *Flaviviridae* also houses viruses known to cause neglected tropical diseases of the genus *Flavivirus*, which comprises over 100 different pathogens including notable viruses such as dengue (DENV), West Nile (WNV), yellow fever (YFV), and zika (ZIKV) (5).

*Flaviviridae* have linear, single-stranded positive RNA genomes of 9–13 kb. These genomes contain a single polyprotein, divided into structural and non-structural genes, and flanked by a 3’ and a 5’ untranslated region (UTR).

The combination of the information from ViralZone (6) and the annotated genomes of *Flaviviridae* from the National Center for Biotechnology Information Reference Sequence Database (NCBI; RefSeq) indicate that a NS5 gene, composed of NS5A and RNA-dependent RNA polymerase (RdRp), is common to the four genera.

However, not all *Flaviviridae* code the same proteins. The genome of *Flavivirus* consists of three structural (C, prM, E), and eight non-structural (NS1, NS2A, NS2B, NS3, NS4A, 2k, NS4B, NS5) proteins. Three of the proteins of *Flaviviridae* are exclusive (prM, NS1, and 2k) to the family (1). *Hepacivirus* contains two envelope proteins (E1 and E2) while *Pestivirus* contains three (Erns, E1, and E2). The E of *Flavivirus* is equivalent to the E1-E2 dimer in the other genera. *Pegivirus* lacks the capsid protein (C) which is present in the other three genera. It is not clear if an additional C might arise in the genus from cleavage of the N-terminus or from an alternative reading frame. Finally, a Npro protease precedes the C in *Pestivirus*.

Given the medical and economic significance of *Flaviviridae*, there is an increasing interest in understanding their genomic structure and evolution as well as to compare known viruses of this family with less known viruses that may become a health concern in the upcoming years. At the time of writing this report, there were 8,633 complete genomes of *Flaviviridae* available in NCBI’s GenBank and RefSeq databases (7). However, only 5,217 (approx. 60%) of these genomes are annotated beyond the polyprotein level. This lack of genome annotation thwarts comparative analyses at the level of individual proteins.

A review of 58 articles on *Flaviviridae* published since 2018 citing records of complete genomes of *Flaviviridae* in GenBank and RefSeq shows that most of these authors analyze the entire polyprotein sequences with no granular annotation or data partitioning of any kind. Authors who annotate genes within the viral polyprotein are scarce and perform the annotations by using a variety of methods. As such, the annotations are absent or do not follow a consistent methodology. For example, Charles et al. (8) identified conserved domains utilizing blastn and blastp (9). Wu et al. (10) deduced polyproteins on pestiviruses by aligning them to other known sequences of the genus and predicted conserved protein domains using either Pfam (11) and InterProScan (12) or the Conserved Domain Database of NCBI (13). As a final example, Wen et al. (14) predicted coding regions using the Predict Protein Server (15).

Our goal of maximizing the number of fully and consistently annotated genomes of *Flaviviridae* allows us to analyze homologous sequences of proteins of interest individually or in combination with other data. A complete gene dataset for *Flaviviridae* will enable phylogenetic analyses to include various genera in a single analysis since we would be able to partition the data taking into account genes that are not ubiquitous in the family. Without sequence annotation to guide data partitioning, the variation of gene content among different genera make it impossible to include all their genomes in the same analysis. Moreover, the alignment of complete polyproteins, even of closely related flaviviruses, can lead to false homology assumptions in which nucleotides of a gene align with nucleotides of another. Therefore, partitioning is key for a group in which phylogenies frequently include only members of a single genus (using no outgroup sequences). In the absence of outgroup sequences, many authors resort to strategies such as midpoint rooting (e.g., 16–18). However, midpoint rooting is often arbitrary, gives a false sense of the polarity of character change in a tree, and creates artificial groups whose monophyly could be contracted had another root been provided.

To overcome these shortcomings, we developed a new approach to annotate genomes from any of the four genera of *Flaviviridae* and evaluated its potential impact on the phylogenetic analyses of these viruses.

It has not escaped our notice that there are authors (e.g., 17, 19, 20) that have expressed concern regarding the combination of different genera or even species of *Flaviviridae* into the same phylogeny fearing that it could lead to long-branch attraction (LBA) events in which the outgroup would be attracted to the largest ingroup branch. Hence, we also test and discuss LBA events that could arise in such a scenario and present an argument in favor of rooting on outgroup sequences instead of applying midpoint rooting in viral phylogenetics.

## New Approaches

Each genomic sequence of *Flaviviridae* was run through a pipeline that started with parallel GeneWise (comes with Wise v2.4.1, 21) and TransDecoder v.3.0.0 (22) analyses. In TransDecoder, we used dedicated databases downloaded from UniProt (23) and blastp (comes with BLAST v2.4.0+, 9) for homology-based annotation. We also employed hmmscan (comes with HMMER v3.1b2, available at http://hmmer.org/) to search the peptides for protein domains using Pfam (11). If predictions from both GeneWise and TransDecoder matched, we pooled the alignments and calculated distance matrices with distmat (comes with EMBOSS v6.6.0, 24). We applied the distance matrices for outlier testing using the Tukey method (25) with the help of homemade R scripts. The Tukey method to search for outliers leverages on the interquartile range and is applicable to most ranges since it is not dependent on distributional assumptions. It also ignores the mean and standard deviation, making it resistant to being influenced by the extreme values in the range. We removed all sequences that were selected as outliers. See figure 1 for a description of the main steps of this pipeline.

**Fig. 1.**
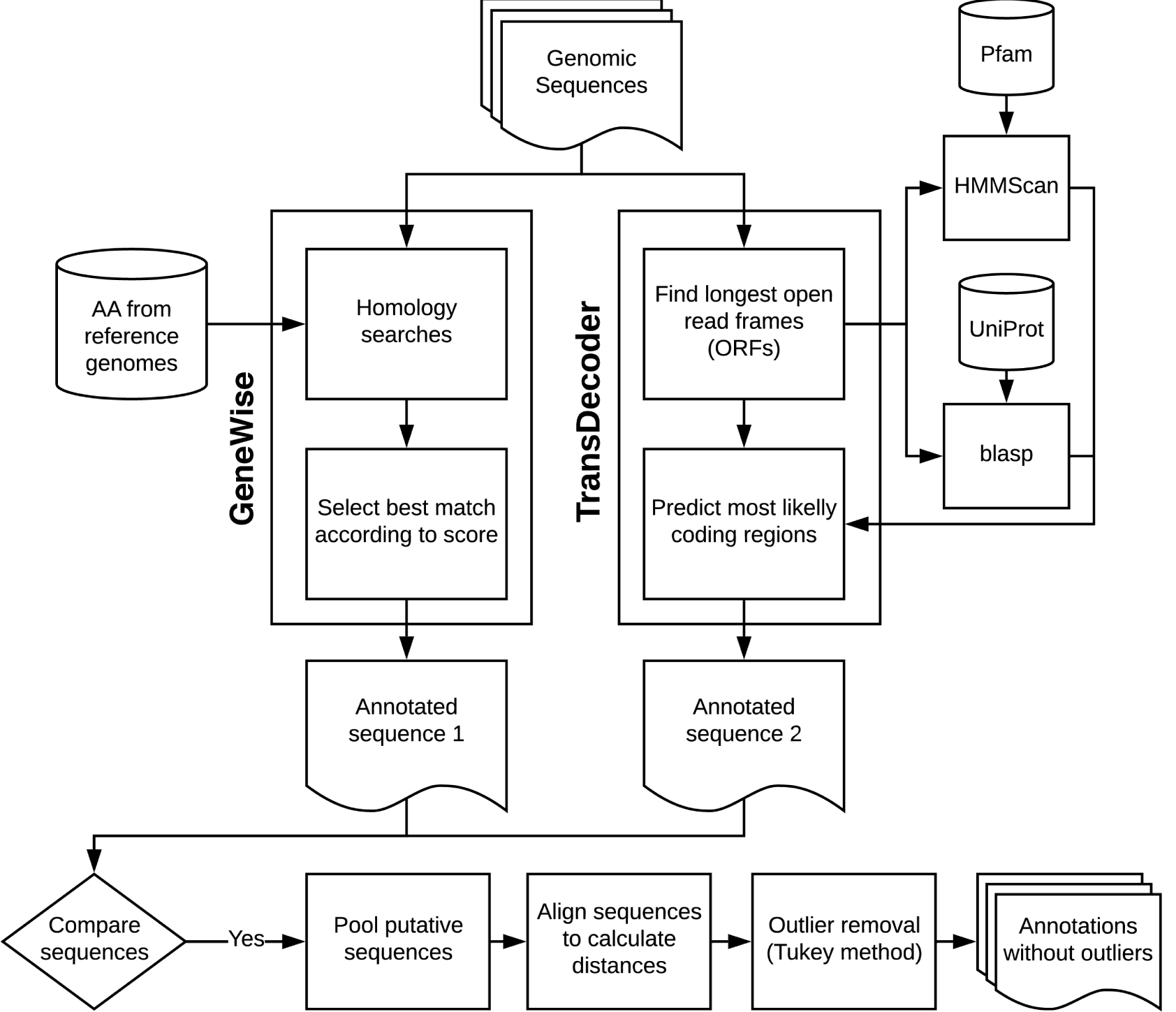
Main steps of the protein annotation pipeline.

## Results

### High efficiency in genome annotation

After the removal of outliers by the Tukey’s method, the pipeline retained 118 complete genomes of *Flaviviridae* (see supplementary table S1, Supplementary Material online). We identified the soybean cyst nematode virus 5 (SbCNV-5) as a single outlier in our dataset (NCBI’s accession number NC_024077). The resulting genomic data comprises 1.25 Mbp (approx. 10,591 bp per genome).

With our pipeline, we re-annotated 63 reference genomes and 55 genomes for which no annotations were available (see supplementary file S1, Supplementary Material online). Although we developed the pipeline on a high-memory server, it is capable of processing the 118 genomes on a personal computer in less than 2 hours. Annotations sum up to 1,247 mature peptides. During manual curation, we were required to perform editing in only 7% of the cases, which corresponded to short fragments such as 2k. Overall annotation efficiency was close to 100%, except for the p7 gene which had an efficiency close to 80%, possibly due to its short sequence length and small prevalence (figure 2).

**Fig. 2.**
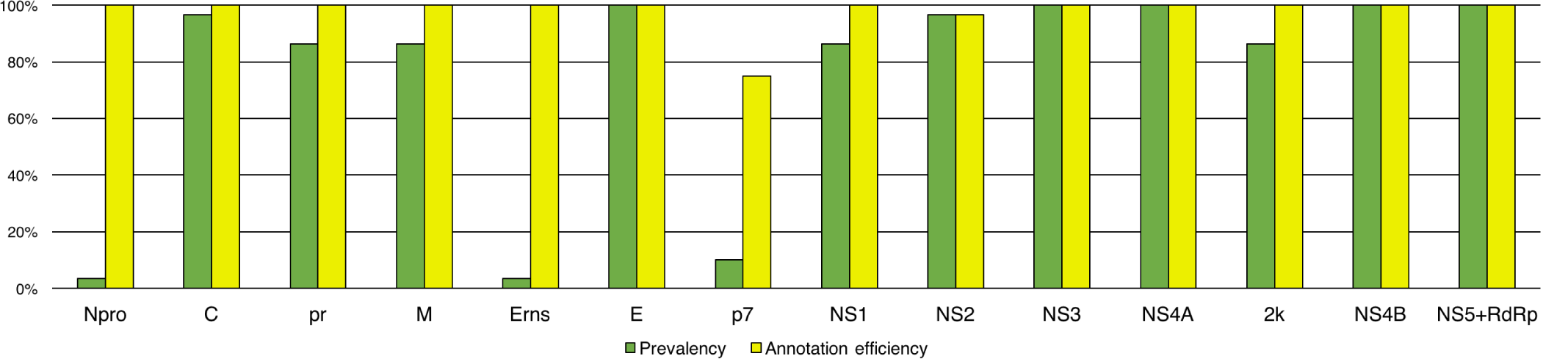
Efficiency of annotation pipeline and partition prevalence.

The pipeline is available at https://gitlab.com/MachadoDJ/FLAVi under a GNU General Public License (GPL) v3.0.

### Topological distances formed two clusters

The matrix of match-split distances among unrooted binary topologies generated hierarchical clusters of trees (figure 3). We did not observe clusters of trees based on optimality criteria or alignment strategy. That is to say that neither the optimality criteria nor the alignment methods seem to strongly direct tree search towards a specific portion of the tree-space.

**Fig. 3.**
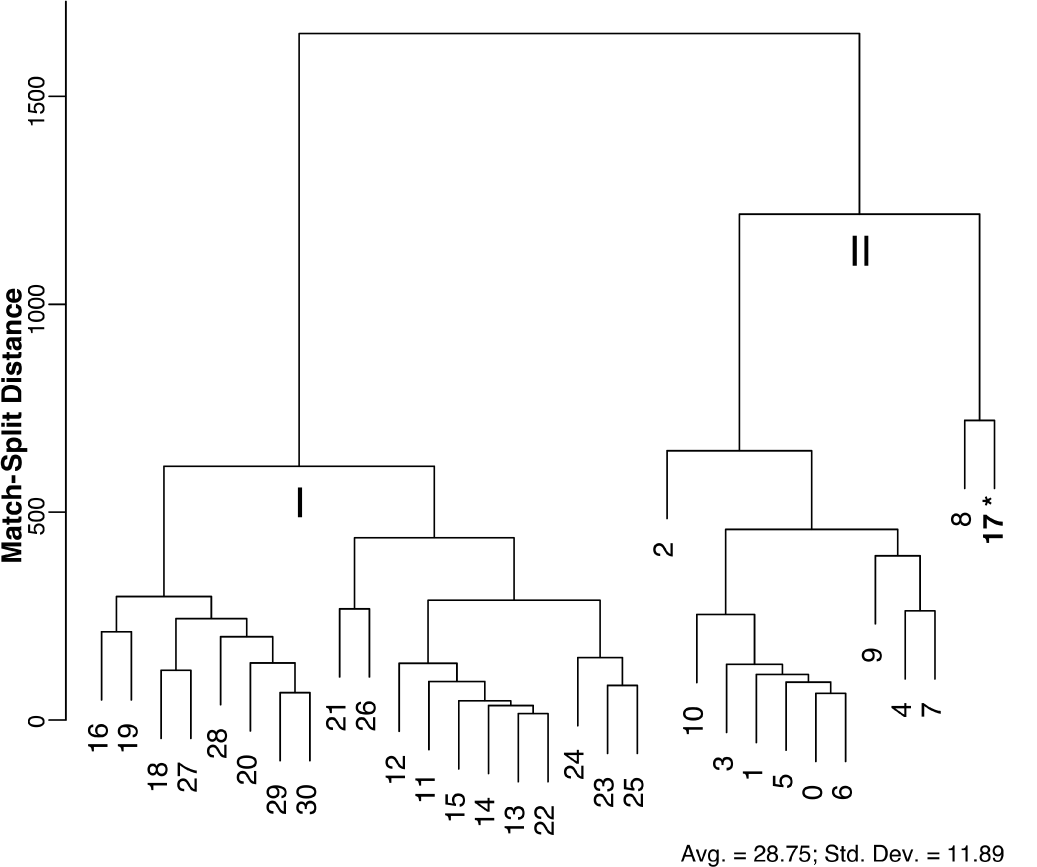
Hierarchical clusters of trees based on match-split distances. Outgroup sequences (*Hepacivirus, Pegivirus*, and *Pestivirus*) were removed to guarantee the compared tree topologies would have the same terminals. Tree numbers correspond to those in table 1. I. No outgroup sequences; some matrices were partitioned. II. Outgroup sequences and partitioned matrices. *This tree was produced without outgroup sequences. See supplementary table S3 (Supplemental Material online) for more details.

However, we can categorize two distant clusters according to the absence (flaviviruses only) or presence (all *Flaviviridae*) of outgroup sequences (groups I and II on figure 3, respectively). The only tree topology obtained without outgroup sequences that was included on group II is a most parsimonious tree (No. 17; see table 1) of flaviviruses, nested beside another most parsimonious tree from the same alignment and optimality criteria.

**Table 1.**
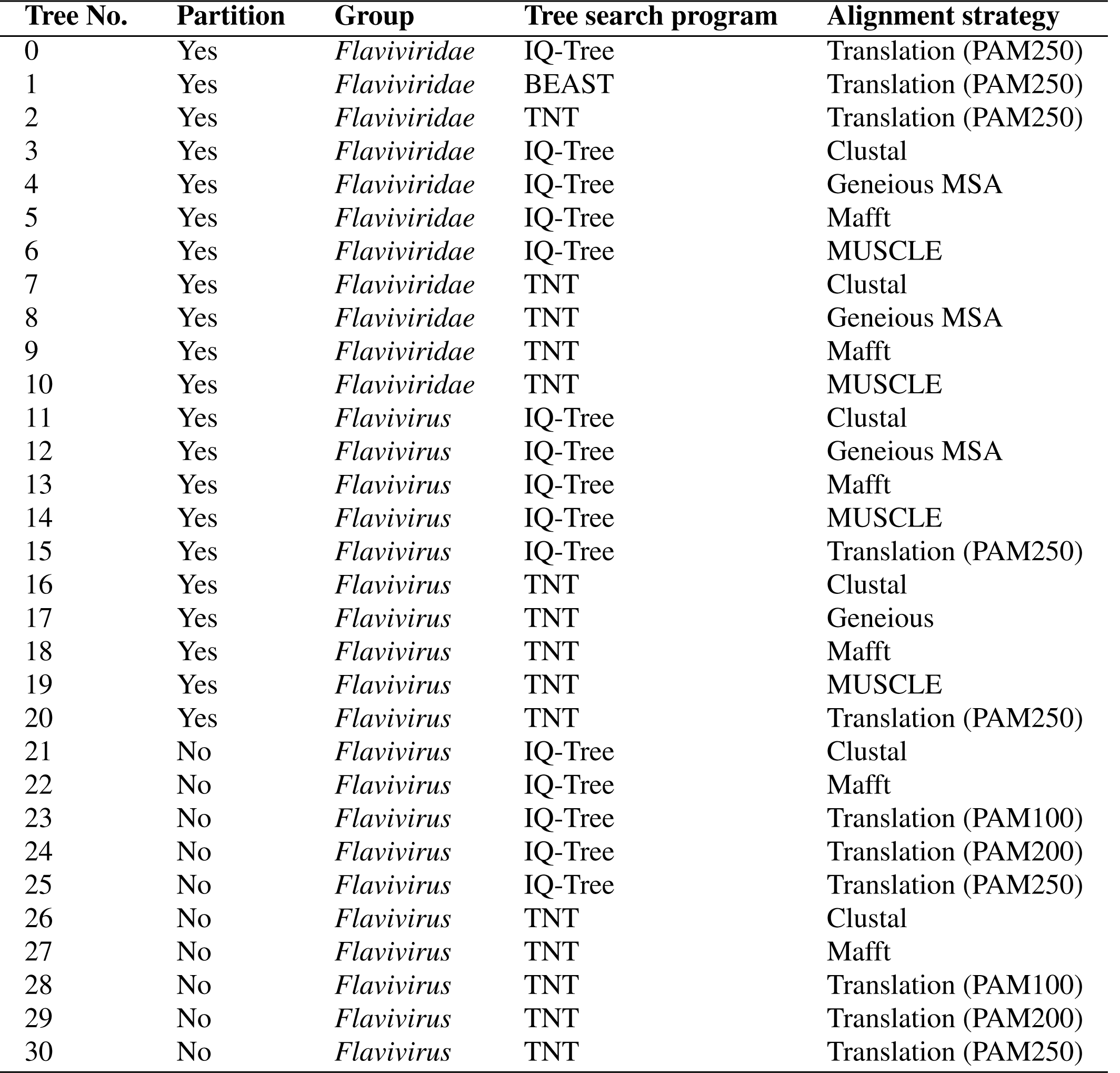
Enumeration of different tree search strategies.

### No long-branch distortions were observed

We compared the topological distances among 11 trees generated during LBA analysis, including three most parsimonious trees (MPTs) produced with total evidence (118 partitioned genomes, aligned using a translation-based method with PAM250 and analysed on TNT) and nine other trees generated when removing each of the six longest terminal branches in our dataset, one at a time: Tamana bat virus (NC_003996; one MPT), simian pegivirus (NC_024377; one MPT), equine pegivirus 1 (NC_020902; one MPT), human pegivirus 2 (NC_027998; three MPTs), rodent pegivirus (NC_021154; one MPT), and Norway rat pegivirus (NC_025679; two MPTs).

Although LBA can affect parsimony (P), we did not observe any two long nonsister branches within clades composed of otherwise short branches in any of the 11 cladograms listed above. We also did not observe any event in which out-group sequences were attracted to the largest ingroup branch (Tamana bat virus).

The calculation of normalized match-split distances shows that the distance among trees from total evidence (original) and trees with one of the long-branches removed (new) vary in similar ways. That is to say that long-branches do not seem to cause any more variation than the variation observed among the most optimal trees.

Kruskal-Wallis test was significant (*p* = 0.0048) only in one of the three comparisons: the variance of distances among new trees and the variance of all trees from the originals (figure 4). This was expected, since our results indicate that the combination of sequence partitioning with an increased sample of outgroup sequences will impact the ingroup relationships. See supplementary file S2 (Supplemental Material online) for details.

**Fig. 4.**
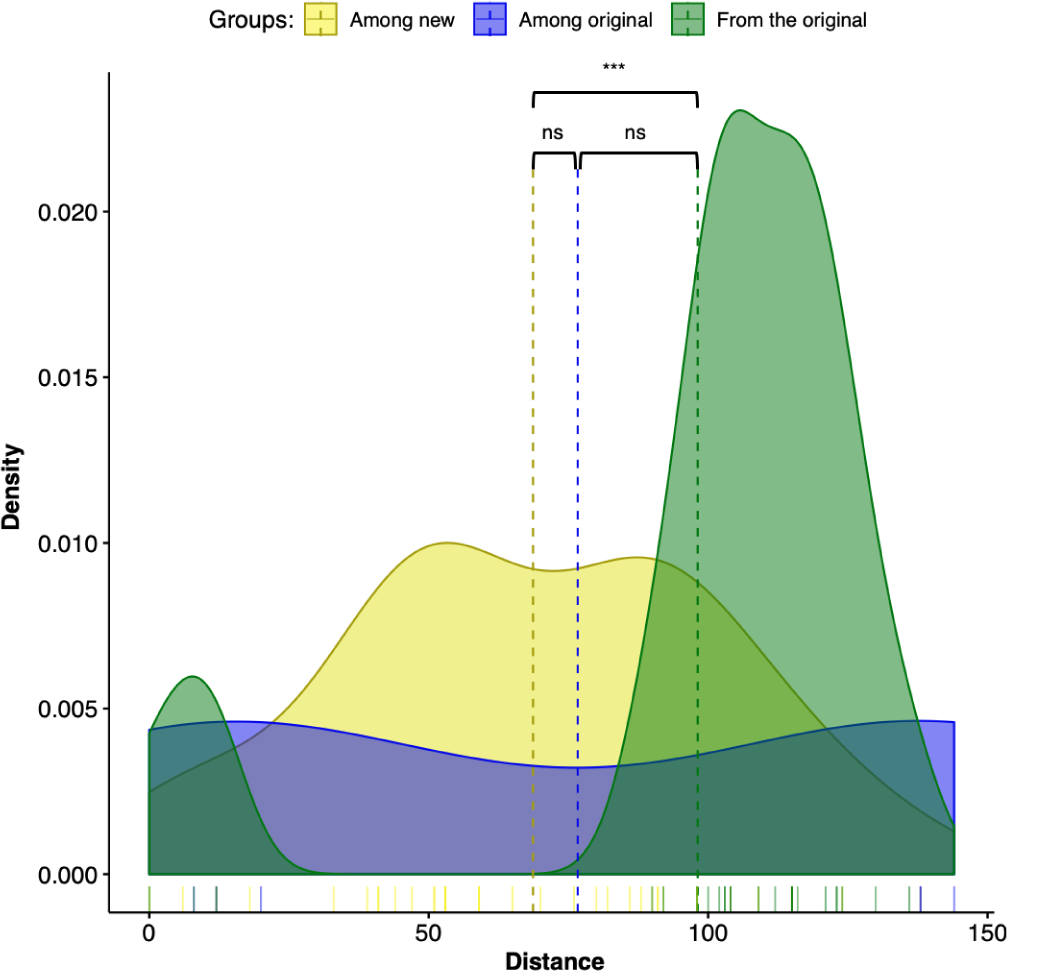
Density plot of the match-split distances among unrooted binary cladograms. Kruskal-Wallis test was significant (*p* = 0.0048) only between groups “among new” and “from the original.” Dashed lines represent the mean of the groups.

### An updated phylogeny of *Flaviviridae*

All 31 tree topologies and 15 matrices generated as a result of our evaluation of the potential impact of tree anno tation and outgroup sampling over tree topology are available in TreeBASE (http://purl.org/phylo/ treebase/phylows/study/TB2:S24096). As the hypothesis, we selected the tree topology from a complete and partitioned matrix aligned with the translation-based method and selected under the maximum likelihood (ML) criteria (tree No. 0 on table 1). We chose ML as our preferred tree-building method given its relatively simple implementation and its ability to incorporate explicit models of molecular evolution while maintaining robustness in the face of differences in base composition and models of nucleotide substitution (26).

We split the evolutionary tree of *Flaviviridae* into two figures. Figure 5 focuses on the genera *Hepacivirus, Pegivirus*, and *Pestivirus* as well as on their relation to *Flavivirus*. Figure 6 presents the relationships among flaviviruses mostly related to tropical diseases transmitted by *Aedes* spp. and *Culex* spp., including DENV, WNV, YFV and ZIKV.

**Fig. 5.**
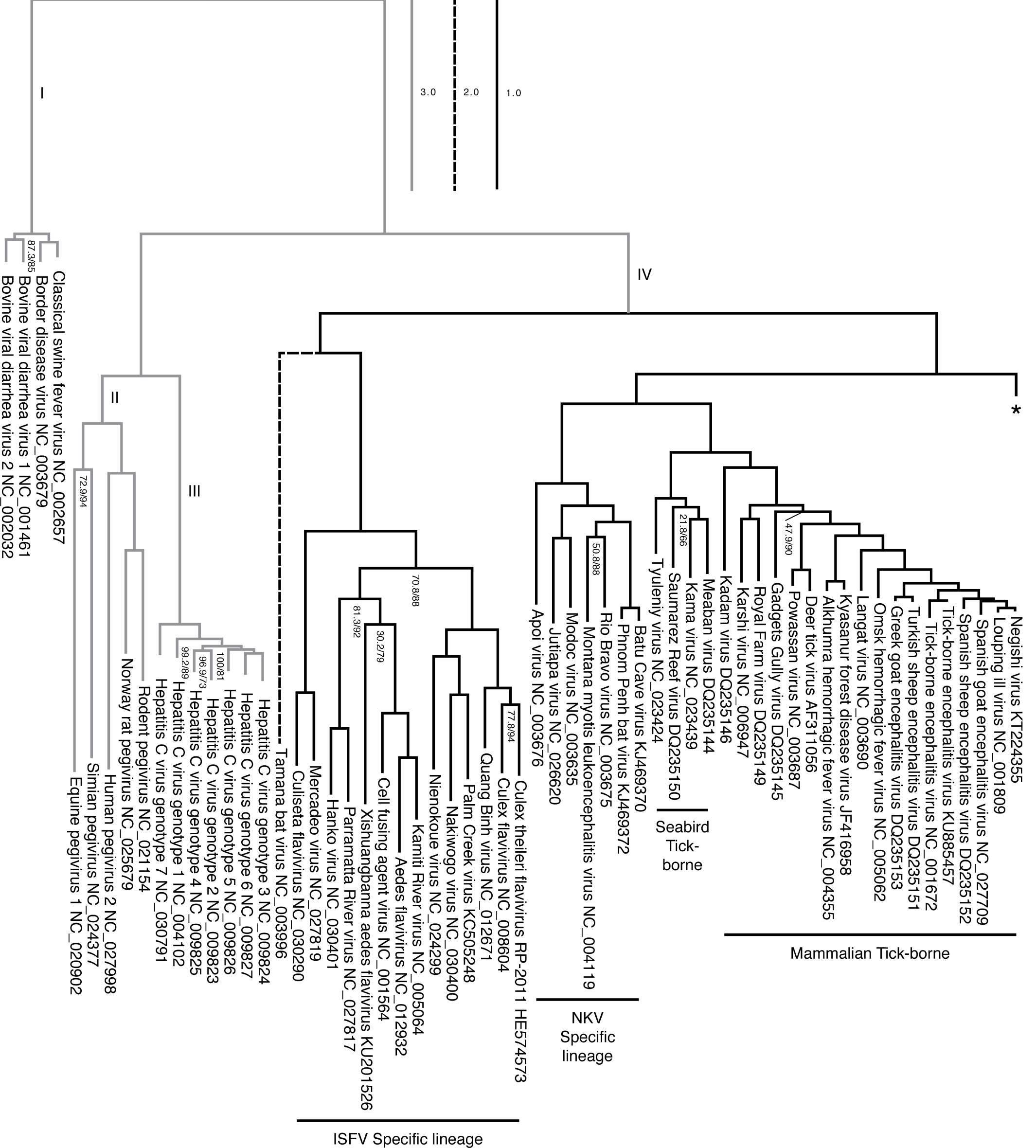
Phylogenetic hypothesis (tree No. 0 in table 1). Branch lengths represent an estimation of the number of substitutions per site. Node labels indicate SH-aLRT support / ultrafast bootstrap (only shown if one of the values is below 90%). Clade names correlate to the character categorization analysis (supplementary file S3, Supplementary Material online). Branch labels represent the four genera: I = *Pestivirus*; II = *Pegivirus*; III = *Hepacivirus*; IV = *Flavivirus*.

**Fig. 6.**
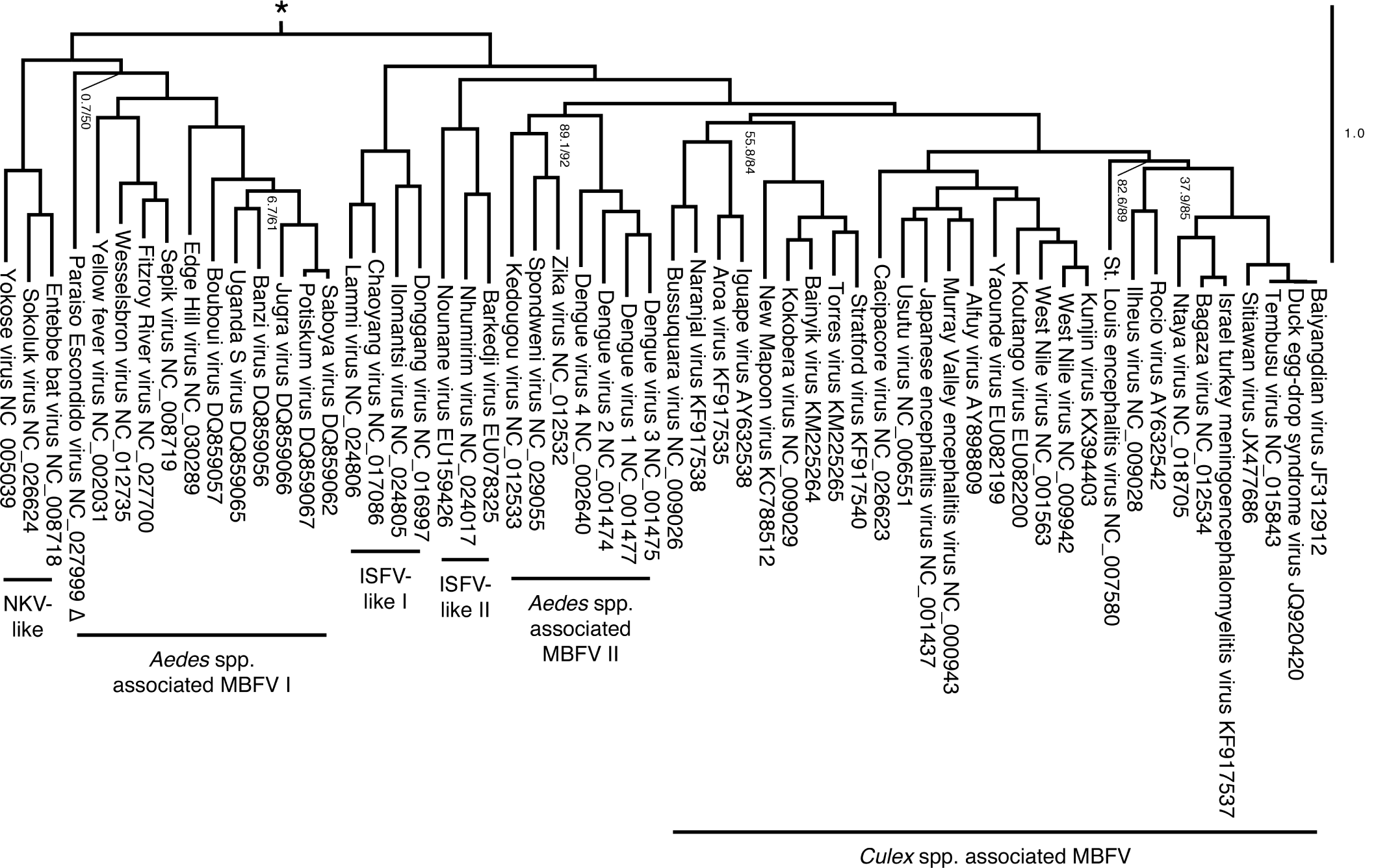
Continuation of the working phylogenetic hypothesis of *Flaviviridae* (see tree No. 0, table 1, figure 5), showing a group within flaviviruses which is sister to (NKV Specific lineage + (Seabird Tick-borne + Mammalian Tick-borne)). Δ = The Ecuador Paraiso Escondido virus (EPEV) was isolated from sand flies (*Psathyromyia abonnenci)*. The EPEV was the first sand fly-borne flavivirus identified in the New World.

Overall, the tree shows high support and values for most clades and corroborated the major groups of *Flavivirus* as presented by Moureau et al. (18). The insect-specific flaviviruses (ISFV-specific lineage), a group that infects only insects, was recovered as sister group to the remaining flaviviruses. All the remaining flaviviruses were found to be the sister group of flaviviruses with no known vector (NKV-specific lineages) and tick-borne flaviviruses (TBFV). The tree also divides all TBFV into two distinct groups of pathogenic flaviviruses of ticks that primarily feed on or parasitize either seabirds or mammals (figure 5).

Similarly to what was observed by Moureau et al. (18), our analysis shows that three flaviviruses (Sokoluk, Entebbe bat, and Yokose virus) appear to have diverged within the mosquito-borne flaviviruses (MBFV) and lost their mosquito association completely, which justify entitling this clade NKV-like group. The NKV-like viruses are sister to the *Aedes* spp. associated MBFV I group, and both comprise a clade that is the sister group to the remaining flaviviruses, including ISFV-like and MBFV clades (figure 6).

Our results also agreed with the phylogenetic analysis of Moureau et al. (18) on finding the ISFV-like lineages I and II to be paraphyletic. The ISFV-like II is the sister group to the clades *Aedes* spp. associated MBFV II and *Culex* spp. associated MBFV (figure 6).

Despite agreeing to the principal clades in Moureau et al. (18) and the relationships among them, our analysis bring additional insights to the phylogenetic relationship within each of them. We draw attention particularly to the internal relations of the clade *Aedes* spp. associated MBFV II in figure 6, which places DENV and ZIKV in a much closer relationship than the one observed in Moureau et al. (18).

## Discussion

### Flexible and conservative genome annotation

We took a conservative approach to annotate the genomic sequences to reduce false positives and minimize improper annotation of highly divergent sequences. The pipeline identified SbCNV-5 as the only outlier in our analyses. Upon careful examination, we observed that the SbCNV-5 genome length is 19.199 bp, approximately 46% larger than the largest genome size expected for *Flaviviridae*. When Bekal et al. (27) described SbCNV-5, they assigned it to *Flavivirus* after presenting protein homology to *Pestivirus*, citing that SbCNV-5 has genome structure, sfRNAs and viral maturation similar to flaviviruses. Nevertheless, the authors mention that SbCNV-5 has an enveloped spherical shaped virion that resembles flaviviruses, but larger (80 nm of size, compared to 50 nm in flaviviruses). The authors explain the difference in size to relate to the larger size of its genome. Although we cannot exclude the hypothesis of SbCNV-5 to share deep ancestry with the current genera of *Flaviviridae* viruses, we cannot place it within flaviviruses or any other of the current genera of *Flaviviridae*. In spite of the fact that we cannot infer the phylogenetic relationships of SbCNV-5 from our analysis due to its high dissimilarity to other members of *Flaviviridae*, this scenario could indicate that SbCNV-5 could land as a new genus of this family having as one of its characteristics a larger genome. It is also possible, however, that SbCNV-5 will become the link to a new family of viruses. More data on relatives of SbCNV-5 will be useful to test these scenarios.

### Outgroup comparison and LBA

Current phylogenetic strategies for genera of *Flaviviridae* often rely on the midpoint rooting of evolutionary trees. Authors are often driven towards midpoint rooting by the perceived difficulty of establishing a correct outgroup to act as root and the possibility of long-branch distortions, especially LBA (17, 28, 29).

The concern about LBA when including outgroup sequences in phylogenetic analysis is justifiable for all optimality criteria in cladistics (parsimony, maximum likelihood, or posterior probabilities calculated for Baeysian inferences). Although LBA was originally described by Felsenstein (30) as a bias in parsimony analysis, the specialized literature have demonstrated that likelihood and Bayesian inference are not immune to it (31). Moreover, we now know that likelihood may also be affected by long-branch distortions, including long-branch repulsion (LBR), in which a true long-branch is not recovered (32). However, the concern with LBA should not impede outgroup comparison when outgroups are available.

Outgroup comparison serves to root the topology and polarize character transformations (33, 34). This comparison is required to convert a network of abstract connections into a concrete evolutionary hypothesis (35). Outgroup comparison also serves as a test of nested hypotheses of ingroup topology and homology (see 36, 37).

Furthermore, empirical studies have demonstrated that increasing outgroup sampling may impact the phylogenetic relationship with ingroup terminals, adding support to clades that otherwise would not be recovered (Grant et al. *et al.* 2017; also see Grant, *submitted manuscript*). Thus, the relevance of outgroup comparison in phylogenetic systematics justifies the effort of dealing with possible long-branch distortions.

Since the specialized literature introduced the first empirical examples of LBA, we have access to different criteria to try to assess whether LBA could have affected the analysis or not (40). These strategies are often based in observing well supported clades of sufficiently long-branches within groups that otherwise include short branches. We made no such observations in our phylogenetic hypothesis. Furthermore, the removal of one long-branch from the tree and the observation of its effect on tree topology could also be an indication of distortions caused by them. Still, pruning long-branches from the trees and calculating the topological distances among them does not show any indication of LBA. Thus, we conclude that LBA did not affect the final phylogenetic hypothesis presented herein and that future research can rely on the same strategies employed here if authors are interested in locating possible LBA in their analysis. Nonetheless, it is worth noting that, since the real history of a group is not known, we cannot guarantee that long-branches do not represent legitimate sister taxa.

As noted by Wheeler (41), if we include a pair of *Panorpa* species into an analysis of insect orders (42), the edge joining these taxa would show a great deal of change. Nevertheless, there would be little reason to doubt such a group.

### Previous work on the Phylogeny of *Flavivirus*

Before 2012, the phylogeny of *Flavivirus* did not include the ISFV-Specific Lineage, and mainly relied on utilizing the Cell Fusing Agent (CFA) as the outgroup of choice given its genetic distance to the other viruses of the genus (43–46). Cook et al. (47) introduced a comprehensive phylogenetic study where it included multiple insect-specific viruses that form together a monophyletic clade, which includes CFA, as a sister group to all other *Flavivirus*. These authors also performed an analysis where they included individual sequences from other genera of *Flaviviridae, Hepacivirus* and *Pestivirus* and find *Hepacivirus* as a sister group to *Flavivirus*, and *Pestivirus* sister to both *Flavivirus* and *Hepacivirus*. Our results are in agreement with these authors in regards to the main clades positioning on the trees, although the increase in the number of taxa has added more information within each vector based group.

Moreover, Zanotto et al. (48) and Twiddy et al. (49) suggested that tick-borne and mosquito-borne flaviviruses have distinct population dynamics. These differences, mainly due to their methods of dispersal, propagation, and changes in the size of the host population, explain the difference on the branching process observed on tick-borne compared to mosquito-borne viruses (figure 5 and 6).

### Novel phylogenetic insights

Multiple researchers evaluated the phylogeny of *Flavivirus* using whole genomes. Although when researchers wanted to increase taxon sampling, the lack of complete genomes available forced researchers to focus on E, NS3, and NS5 genes (43, 45). More recently, multiple groups attempted to generate a phylogeny of *Flavivirus* using the whole polyprotein information (18, 50–52). Our results are in agreement with most studies, except Li et al. (52). Li et al. (52) attempted an alignment-free method based on natural vectors and by doing so have found the tick-borne group sister to all other groups, including the ISFV-specific lineage. These authors present an alternative tree based on multiple sequence alignment of the whole polyprotein and tree building using neighbor-joining, where they could recover ISFV-specific lineage as the sister group to all other flaviviruses, but by doing so, they also find Tamana bat virus (TABV) as a distant sister-group to all flaviviruses.

Appropriate genome annotation allows the alignment of specific gene partitions, making it feasible to deal with variations in gene content and different levels of sequence divergence. Thus, we are capable of including more information into a phylogenetic framework that maximizes the explanatory power of the analysis and allows us to focus not only on the relationships within genera but also relationships between genera. When aligning whole polyproteins, multiple researchers left out the concept that each protein within a polyprotein is a modular unit. In doing so, the current phylogenetic studies assume homology of the whole polyprotein (50, 52), although still assuming homology some take post alignment processing steps for cleaning spurious regions before the phylogenetic analysis (18, 51). Ignoring the homology of individual proteins leads to the generation of spurious alignments and errors which will be perpetuated on down-stream analysis (53).

Our results add new sequences to the *Flaviviridae* phylogeny. The addition of new sequences to the phylogeny based on homology statements may assist on the taxonomy and official classification of these new viruses as part of *Flaviviridae* by groups such as the International Committee on Taxonomy of Viruses.

For instance, similarly to the SbCNV-5, the Tamana bat virus (TABV, NC_003996) is also considered highly divergent from most flaviviruses. De Lamballerie et al. (54) mentions that they could not place TABV in a precise region on the tree, possibly due to the significant genetic differences from other members of the family. Thus, the authors suggested assigning TABV as a new genus. Moureau et al. (18) also considered TABV too divergent to be included given their particular methodology. Due to the partitioning of the data set according to genes annotated with our pipeline, we are capable of including TABV into a phylogenetic framework that shows that, despite its accumulation of genetic transformations, places it with high support within *Flavivirus* as the sister group of the ISFV clade.

Within *Flavivirus*, we also observe that DENV forms a monophyletic clade with ZIKV, Kedougou and Spondweni viruses, all of which are transmitted by *Aedes* spp. mosquitoes. Previously, Moureau et al. (18) identified these three viruses in a sister group to DENV, which encompass the *Culex* spp. associated group and previous related research suggested that the positioning of these viruses was ambigu ous (55). Nevertheless, individual studies focusing on specific viruses have observed this relationship (56). Our results not only add new sequences to the *Flaviviridae* phylogeny, but while doing so, agree with the current literature where we were able to recover the vast majority of known clades, sharing vector specificity. These results serve to bring additional details into the evolution of specific lineages of flaviviruses at the same time they demonstrate the ability of the methods presented here to deal with the diversity of genomes of *Flaviviridae*, including highly divergent sequences.

## Conclusions

The new approach presented here is a dedicated annotation pipeline for *Flaviviridae*. Our annotation pipeline uses a combination of *ab initio* and homology-based strategies and recovered 100% of the genes in reference genomes and showed an annotated over 97% of the expected genes in the remaining sequences that were not curated previously.

The annotation of the genomes of *Flaviviridae* allows the combined phylogenetic analysis of all genera of the family (*Flavivirus, Hepacivirus, Pegivirus*, and *Pestivirus*). The phylogenetic analysis of *Flavivirus* was enhanced for all cladistic optimality criteria (parsimony, maximum likelihood, and posterior probability) when we included outgroup sequences from other genera of *Flaviviridae*. The inclusion of outgroup sequences did not result in any noticeable long-branch distortions.

The final tree sheds light on the phylogenetic relationship of viruses that are divergent from most *Flaviviridae*. In particular, our results indicate that the soybean cyst nematode virus 5 (SbCNV-5) may not be a member of *Flaviviridae*, and that the Tamana bat virus belongs to *Flavivirus* and is especially closely related to insect-specific flaviviruses (ISFV-specific lineage). We also corroborate the close phylogenetic relationship of DENV and ZIKV with an unprecedented degree of support. The main phylogenetic insights of our phylogeny reconstruction were made possible due to the inclusion of all genera of *Flaviviridae* into a single analysis which maximized the explanatory power of the data and enhanced the resolution of phylogenetic relationships in the ingroup.

## Materials and Methods

### Taxon sampling

We downloaded a total of 78 complete reference genome sequences of *Flaviviridae* from NCBI’s Reference Sequence (RefSeq) database (https://www.ncbi.nlm.nih.gov/refseq/). We downloaded 41 complete genome sequences of *Flaviviridae* from NCBI’s GenBank (https://www.ncbi.nlm.nih.gov/genbank/). These sequences comprise four genomes of *Pestivirus*, five genomes of *Pegivirus*, seven genomes of *Hepacivirus*, and 103 genomes of *Flavivirus*, for a total 119 genomes. The total nucleotide count of the data is Mbp or 10,663 bp on average per genome. Of all these genomes, 63 were already annotated. See supplementary table S1, Supplementary Material online.

We selected *Flavivirus* as ingroup since this genus has the highest diversity within our taxon sampling and therefore is more likely to provide insights on changes of topology due to the influence of outgroup selection and data partitioning.

### Protein partitions

Combined, we can divide the polyproteins of the four genera of *Flaviviridae* into 14 protein regions, although not all regions are common to the entire family (figure 7). To understand the effects of data partitioning and outgroup sampling on the phylogenetic relationships of *Flaviviridae*, we used three different strategies to partition the data for the phylogenetic analysis based on genome annotation. First, we used single partition for the entire coding region for each polyprotein in *Flavivirus*. Second, we used 11 partitions for the analysis of sequences of *Flavivirus* (i.e. all partitions except Npro, Erns, and p7). Lastly, we used 14 partitions including all four genera, in which we coded absent genes as missing data.

**Fig. 7.**
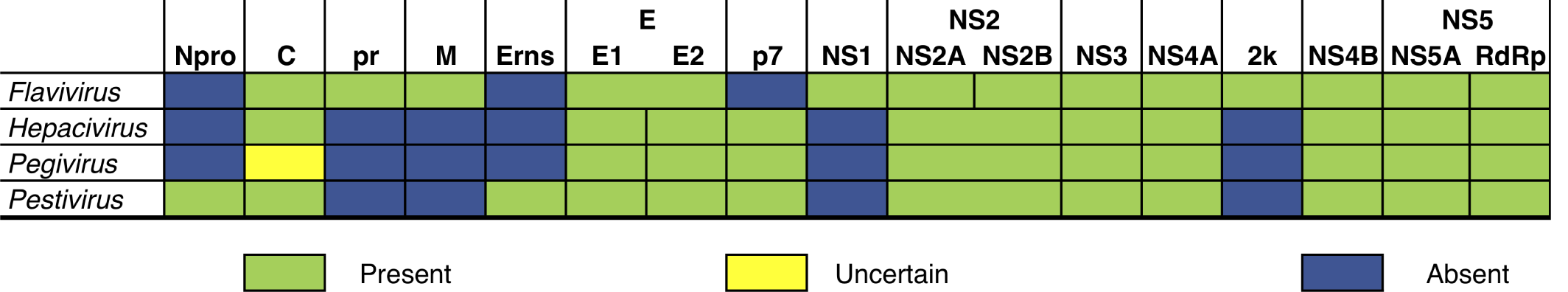
Genome structure comparison between the four genera of *Flaviviridae* (*Flavivirus, Hepacivirus, Pegivirus* and *Pestivirus*), showing the 14 partitions that can be found in the genomes of the family.

### Tree search

We applied the following multiple sequence alignment techniques to account for the sensitivity of tree topology to the variation on homology statements at the character level: MAFFT, Clustal, MUSCLE, and translation based alignment under different scoring matrices (PAM100, PAM200, and PAM250). We used Geneious (v8.1.9) (57) to execute every alignment using default arguments. Our criteria for selection of best alignment strategies was the maximization of overall sequence identity and minimization of alignment length.

To focus on the effects of data partitioning and to minimize possible artifact caused by missing data on partitions of proteins that are not found in all *Flaviviridae*, we built some datasets with 1) complete polyproteins and 2) partitioned proteins of flaviviruses that contained all partitions. To address the sensitivity of tree topology to the inclusion of outgroup sequences, we used partitioned data on analyses containing either exclusively flaviviruses or all the four genera of its family. Finally, since tree search can be affected by the choice of optimality criterion, we explored the results of three phylogenetic tree search methodologies: TNT 64-bit version with notaxon limit (parsimony – P) (58), IQ-Tree v1.6.1 (maximum likelihood – ML) (59) and BEAST v2.4.8 (Bayesian inference – BI, under the criterion of posterior probability – PP) (60). See supplementary table S2, Supplementary Material online, for non-default arguments used for each tree search. In total, we performed 31 tree search analyses considering different alignments and optimality criteria. Of those, ten trees are from whole polyprotein alignments of flaviviruses, ten are from the partitioned datasets of flaviviruses, and 11 are from the partitioned datasets of the combined four genera of *Flaviviridae*, as shown in table 1. We selected the ML tree from a complete and partitioned matrix aligned with translation-based method as our hypothesis, but we made all matrices and tree search results available in TreeBase (https://www.treebase.org, see Results section). For the final hypothesis from ML, we executed the SH-aLRT measure of support (61) and ultrafast bootstrap analysis of clade frequencies (62). The SH-aLRT is an approximation of the likelihood ratio, a direct measure of how much the evidence supports the hypothesis. SH-aLRT is an approximation of the ratio of the log-likelihood of the optimal hypothesis and the best contradictory hypothesis. The ultrafast bootstrap analysis is a variation of the traditional bootstrap using heuristics and constraints to speed the search for optimal tree topologies, and the specialized literature shows that it is largely consistent with the traditional bootstrap analysis (62).

### Sensitivity analysis

We pooled all 31 cladograms, pruning off outgroup branches when necessary, to calculate the pair-wise distance matrix among the phylogenetic hypothesis of flaviviruses based on match-split distances (MSdist; 63) with the program MSdist v1.0. The match-split distances focus on the topological distance of unrooted phylogenetic trees based on splits, similarly to the Robinson-Foulds metric (RF; 64). However, it is more sensitive and resistant than RF and to single terminal displacements. A homemade R script (65) was used to create a dendrogram and distance plot summarizing the distances among all tree topologies.

### Long-branch attraction analysis

Felsenstein (30) introduced the issue of long-branch attraction (LBA) in phylogenetics as a problem of statistical inconsistency that affects parsimony (P). Felsenstein (30) advocated maximum likelihood (ML) as an alternative. The issue becomes apparent “when two nonsister branches are long while other branches are short” (see 66, p. 202, for a historical perspective). However, there are also numerous accounts of matters of similar nature (e.g., long-branch repulsion) affecting ML, showing that it can also be inconsistent under certain conditions (e.g., 67–69).

Authors such as Thézé et al. (17, p. 2998) have raised concerns on possible effects of long-branches from the outgroup, choosing to avoid the inclusion of terminals outside the in-group altogether: “Midpoint rooting was chosen to root ML trees in order to avoid long-branch attraction with highly divergent outgroups.” Therefore, we evaluated the potential influences of long-branches on the hypothesis where we included the four genera of *Flaviviridae*.

We selected six terminals on our final topology that were visibly longer than all others and removed them, one at a time, from the complete dataset. This resulted in six additional matrices that we analyzed with TNT since P is more likely than ML to be affected by LBA. We used a homemade Python v3.6 script to prune all these terminals from every cladogram and applied MSdist to calculate the distance among all trees. Finally, we performed the Kruskall-Wallis test to evaluate the differences in the variance among different groups of trees.

### Character categorization

We carried the categorization of synapomorphies with YBYRÁ (70) to evaluate important mutations on clades of particular epidemiological interest. YBYRÁ indicates if a derived state occurs only in the clade in question (non-homoplastic) or also occurs in other clades (homoplastic) and if it is shared by all terminals of the clade (unique) or is subsequently transformed into one or more different states within the clade (non-unique). Character categorization is useful to both support and describe clades.

### Computational resources

We performed all annotation and tree search in Heket, a high-memory server housed in the Museum of Zoology of the University of São Paulo (see http://www.ib.usp.br/grant/anfibios/researchHPC.html). Heket has a dual Processor Intel Xeon E52620v2 (24 cores), 256GB DDR3 ECC, 4 x HDD 4Tb (10,8Tb RAID 5.0), SSD 240GB, and Infini-band (20 Gb/s). It runs on an Ubuntu Linux v16.04 OS. We used a MacBook Pro (macOS High Sierra 10.13, 2.4GHz Intel Core i5, 16 GB RAM) for alignment manipulation and analysis of results.

## Supporting information

configuration.xml

README

GenBank.gb

RefSeq.gb

## Supplementary Material

Supplementary digital material, including tables S1–4 and files S1–3, is available at bioRxiv online (https://www.biorxiv.org/).

## Acknowledgements

We acknowledge the Department of Bioinformatics and Genomics, the College of Computing and Informatics, the Ribarsky Center for Visual Analytics, and the Graduate School of The University of North Carolina at Charlotte. We are thankful for the computational resources made available to us by Taran Grant from the Laboratório de Anfíbios Zoology Department of the University of São Paulo, Brazil (FAPESP Proc. No. 2012/10000-5). We also thank Michael T. Wolfinger for comments on the early version of this manuscript. This work was partially supported by the National Institutes of Health (NIH) National Institute of Allergy and Infectious Diseases (grant number AI135992) to ABS.

